# Intranasal VLP-RBD vaccine adjuvanted with BECC470 confers immunity against Delta SARS-CoV-2 challenge in K18-hACE2-mice

**DOI:** 10.1101/2023.04.25.538294

**Authors:** Katherine S Lee, Nathaniel A Rader, Olivia A Miller, Melissa Cooper, Ting Y Wong, Md. Shahrier Amin, Mariette Barbier, Justin R Bevere, Robert K Ernst, F. Heath Damron

## Abstract

As the COVID-19 pandemic transitions to endemic, seasonal boosters are a plausible reality across the globe. We hypothesize that intranasal vaccines can provide better protection against asymptomatic infections and more transmissible variants of SARS-CoV-2. To formulate a protective intranasal vaccine, we utilized a VLP-based platform. Hepatitis B surface antigen-based virus like particles (VLP) linked with receptor binding domain (RBD) antigen were paired with the TLR4-based agonist adjuvant, BECC 470. K18-hACE2 mice were primed and boosted at four-week intervals with either VLP-RBD-BECC or mRNA-1273. Both VLP-RBD-BECC and mRNA-1273 vaccination resulted in production of RBD-specific IgA antibodies in serum. RBD-specific IgA was also detected in the nasal wash and lung supernatants and were highest in VLP-RBD-BECC vaccinated mice. Interestingly, VLP-RBD-BECC vaccinated mice showed slightly lower levels of pre-challenge IgG responses, decreased RBD-ACE2 binding inhibition, and lower neutralizing activity *in vitro* than mRNA-1273 vaccinated mice. Both VLP-RBD-BECC and mRNA-1273 vaccinated mice were protected against challenge with a lethal dose of Delta variant SARS-CoV-2. Both vaccines limited viral replication and viral RNA burden in the lungs of mice. CXCL10 is a biomarker of severe SARS-CoV-2 infection and we observed both vaccines limited expression of serum and lung CXCL10. Strikingly, VLP-RBD-BECC when administered intranasally, limited lung inflammation at early timepoints that mRNA-1273 vaccination did not. VLP-RBD-BECC immunization elicited antibodies that do recognize SARS-CoV-2 Omicron variant. However, VLP-RBD-BECC immunized mice were protected from Omicron challenge with low viral burden. Conversely, mRNA-1273 immunized mice had low to no detectable virus in the lungs at day 2. Together, these data suggest that VLP-based vaccines paired with BECC adjuvant can be used to induce protective mucosal and systemic responses against SARS- CoV-2.

## Introduction

Vaccines prepare the immune system to defend against disease-causing agents through a simulated exposure. Attenuated or inactivated whole-viruses, viral vectors, pathogen proteins, mRNA, and bacterial toxoids, are all used in vaccine formulas administered to hosts to train the immune system to adequately fight off pathogens without an authentic exposure or infection. The immunological memory response that develops from this priming exposure event is oftentimes highly capable of protecting against severe disease and death. Still, the vaccine-elicited immune response differs from a natural response to pathogen exposure and may not always produce superior cellular responses. The intramuscular route of administration is a popular option for vaccine delivery strategies due to its low incidence of site-specific adverse reactions, and optimal immunogenicity for a systemic immune response [1]. Blood supply to the muscles, compared to other tissues, clears adjuvants and other vaccine components more rapidly, leading to the efficient uptake and spread of antigen through the periphery. Circulating antigens are then directed to the lymph nodes where they come into contact with antigen presenting cells (APC) in germinal centers for the generation of antigen-specific T cell and B cell responses [2].

Despite the ability of intramuscular vaccines to induce systemic protection against specific pathogens, alternate routes of administration have been investigated to improve comfort and boost the timeframe of protection through elicitation of site-specific immune responses [3,4]. Intranasal vaccination is an attractive delivery route for its ability to recapitulate natural infection by respiratory pathogens, priming the mucosa—a site otherwise difficult to generate adaptive immunity in—to provide protection when its needed [5]. There is only one intranasal vaccine currently approved for human use worldwide—FluMist Quadrivalent—although its use is restricted to healthy, non-pregnant individuals between the ages of 2 and 49 due to concerns surrounding its immunogenicity and the use of a live-attenuated virus [6]. With eight orally administered vaccines (the United States have utilized oral vaccines against rotavirus, polio, and others), this together makes a limited number of whole-virus vaccines that directly target the mucosal arm of the immune system [7]. We, and others, hypothesize that a protective mucosal immune response may be the key to ultimate protection against pathogens including SARS-CoV-2 that primarily target and replicate in the upper respiratory tract’s mucosal surfaces.

Intranasal COVID-19 vaccines are highly desirable as the COVID-19 pandemic evolves and persists [5,8,9]. Not only could they circumvent the discomfort occurring after intramuscular vaccination using the current mRNA-based COVID-19 vaccines, but they may also induce a superior level of protection. In the case of COVID-19 vaccines, studies have shown that vaccinated hosts have relatively weak neutralizing antibody responses against multiple SARS-CoV-2 variants compared to convalescent patients [10]. Specifically, the utilization of intramuscular mRNA vaccines now and into the future as seasonal boosters poses an even bigger problem as it does little to induce mucosal respiratory immunity that is integral to stopping viral replication and therefore preventing high transmission rates among hosts infected with transmissible variants like Omicron. India and China were the first countries to approve the vaccines BBV154 and Ad5-nCoV-S (respectively) for intranasal use [11]. By vaccinating intranasally from the beginning, or by introducing intranasal boosters on top of a completed intramuscular vaccine schedule, the primed immune response against SARS-CoV-2 exposures can evolve to include pathogen specific IgA antibodies as well as greater B cell and tissue-resident memory (T_RM_) responses throughout the respiratory tract [12–15].

Previously, our lab developed the experimental BReC-CoV-2 vaccine that was effective against SARS-CoV-2 challenge in K18-hACE2-transgenic mice when two doses were administered either first intramuscularly (prime) then intranasally (boost), or both intranasally [16]. To build upon the protection of this formulation when administered intranasally, we aimed to improve the overall immunogenicity by utilizing a virus-like particle (VLP) antigen containing SARS-CoV-2 WA-1 Receptor Binding Domain (RBD) proteins conjugated using the SpyTag system to a Hepatitis B surface antigen (HbsAg) [17]. The VLP-RBD particle increases the ratio of RBD antigen in the vaccine formulation compared to the RBD-CRM used in BReC-CoV-2. Similar to BReC-CoV-2, this VLP-based vaccine was then adjuvanted with the TLR4-agonist Bacterial Enzymatic Combinatorial Chemistry 470 (BECC 470) to enhance both the cellular and humoral immune responses [18,19]. Intranasal administration of two doses of the VLP-RBD-BECC vaccine to K18-hACE2 mice provided equal protection against disease manifestation and morbidity, as compared to two intramuscular administrations of an mRNA vaccine (mRNA-1273; 1/10^th^ human dose) after intranasal challenge with a lethal dose of the SARS-CoV-2 Delta variant. VLP-RBD-BECC limited viral replication in the upper airway at two days post-challenge and maintained a reduction in viral RNA in the nasal wash, lung, and brain, between days two and 10. VLP-RBD-BECC vaccinated mice showed similar serum IgG titers to mRNA vaccinated mice. Importantly, intranasal VLP-RBD-BECC elicited greater RBD specific IgA antibody levels in the lung and nasal cavity than mRNA vaccination or challenge alone. Compared to mRNA, VLP-RBD-BECC consistently maintained low histopathological inflammation scores and concentrations of proinflammatory CXCL10 in the lung tissue as well. VLP-RBD-BECC elicited reduced amounts of Omicron-specific antibodies, however, two doses of the vaccine still demonstrated some protection against viral replication in the lungs of mice following challenge with the Omicron variant.

## Methods

### Ethics and Biosafety Statement

B6.Cg-Tg(K18-ACE2)2Prlmn/J transgenic mice were purchased from Jackson Laboratories and used for vaccination and viral challenge studies under West Virginia University IACUC protocol #2009036460. Mice were continuously monitored for adverse reactions to vaccination and for morbidity after challenge and were humanely euthanized according to our lab’s disease scoring system. West Virginia University’s Biosafety Level 3 Laboratory was used for SARS-CoV-2 challenge studies under IBC protocol #20-09-03. Before additional analysis in BSL2 laboratory space, all mouse tissues obtained from BSL3 were treated with 1% Triton by volume or placed in TRIzol (Zymo R2050-1) to inactivate virus.

### Formulation of VLP-RBD-BECC vaccine and K18-hACE2 mouse vaccination

SARS-CoV-2 Wu RBD proteins were cloned into and purified from *Komgataella phaffi* as previously described [20–22]. To form the VLP, RBD-SpyTag antigens were incubated overnight with HBsAg-SpyCatcher VLP [23,24]. The BECC 470 adjuvant was obtained from Dr. Robert Ernst at the University of Maryland [18]. Vaccines were prepared in batch by sonicating 25µg BECC 470 per dose in water for 15min, then adding RBD-HBsAg VLP (10µg per dose) and incubated rotating at room temperature for 2hrs. Before administration, 10X PBS was added to bring the dose volume to 50µL. Female 7-week-old K18-hACE2 mice were intranasally vaccinated with 25µL per nare under anesthesia with intraperitoneal ketamine (Patterson Veterinary 07-803-6637)/xylazine (Patterson Veterinary 07-808-1947). No-vaccine no-challenge control mice were administered 50µL 1X PBS intramuscular in the hind flank. mRNA control mice were administered 50µL mRNA-1273 vaccine intramuscular in the hind flank as well. All vaccine groups received a second identical dose 4 weeks later.

### Quantification of anti-SARS-CoV-2 RBD IgG antibodies by ELISA

Submandibular bleeds to collect serum were performed to assess immunogenicity 4 weeks after prime and boost doses. Serum was also collected at euthanasia via cardiac puncture. Anti-SARS-CoV-2 RBD IgG levels were quantified using ELISA and a method described previously [25]. High binding plates were coated overnight with 2 µg/mL RBD. The next day, plates were washed 3x and blocked with 3% nonfat milk in PBS-Tween20. After an hour incubation at room temperature, plates were washed 3x then prepared for sample: 5 µL of serum in 95 µL 1% nonfat milk-PBS-Tween20 was added to the top row, and 50 µL of 1% nonfat milk-PBS- Tween20 was added to the remaining wells for dilution across two plates. Samples were diluted 1:2 from row A of plate 1 to row G of plate 2, discarding before dilution into row H. Samples incubated shaking at room temperature for 1 hr. Plates were washed 4x before adding 100 µL secondary antibody (goat anti-mouse IgG HRP; Novus Biologicals NB7539) (diluted 1:2000) in 1% nonfat milk-PBS-Tween20 to all wells and shaking for an additional 1 hr at room temperature. Non-bound antibodies were washed with 5x washes and 100 µL TMB substrate was added to all wells. After 15 min incubation in the dark, 25 µL 2N sulfuric acid was added to stop development and plate absorbances were read at 450 nm on the Synergy H1 plate reader. Serum antibody levels were quantified using Area Under the Curve analysis in GraphPad Prism V9.0.0.

### Quantification of anti-SARS-CoV-2 RBD IgG antibodies by ELISA

To measure anti-RBD IgA in K18-hACE2 mouse tissues following challenge, the ELISA protocol described previously was utilized with minor adaptations. High-binding 96-well plates were coated with RBD, washed and blocked following the same protocol as was used to measure IgG. Mouse samples were added to the top row of wells (5 µL of serum in 95 µL 1% nonfat milk-PBS-Tween20; 20 µL lung supernatant in 80 µL 1% nonfat milk-PBS-Tween20; 100 µL nasal wash), then diluted down the columns 1:2 discarding before the last well. After 2 hours of incubation shaking at room temperature, 100 µL anti-IgA secondary antibody diluted 1:10000 (goat anti-mouse IgA HRP; Novus Biologicals NB7504) was added to all wells. Secondary antibody was left to incubate for 2 hours, then plates were washed, developed, and read as previously described.

### In vitro SARS-CoV-2 RBD ACE2 binding assay

Neutralizing potential of serum antibodies collected from vaccinated K18-hACE2 mice pre- and post-challenge was analyzed using MSD’s V-PLEX SARS-CoV-2 Panel 22 Mouse IgG kit (K15563U-2). Binding to the following antigens was assessed: COV-2 RBD, Delta RBD, Gamma RBD, Beta RBD, Alpha RBD, and Omicron RBD. Serum from mice euthanized at 10 days post- challenge or from 4-week post-boost submandibular bleeds was diluted at 1:5, 1:50, 1:500, and 1:5000 and analyzed following the manufacturer’s protocol. Binding (% neutralization) was measured via electrochemiluminescence values compared to baseline wells with no antibody binding. The percent neutralization 50 (PRNT50) was calculated by fitting a nonlinear regression curve to the plotted percent binding values and interpolating unknowns in GraphPad Prism version 9.

### In vitro authentic SARS-CoV-2 plaque reduction assay

Vero E6 ACE2/TMPRSS2 cells were plated at 70,000 cells per well in 24-well plates and incubated at 37°C and 5% CO2 for 24 hours. Stock SARS-CoV-2 Delta virus (B.1.617.2 hCoV- 19/USA/WV- WVU-WV118685/2021) was diluted in supplemented DMEM media. Mouse serum was diluted 1:5 then ten-fold serial dilutions in media. Diluted mouse serum was mixed with diluted virus in a 1:1 (v/v) ratio and incubated at room temperature for 30 minutes. Cell media was aspirated. 100μl of sample (serum + virus) was added to each well. Plates were incubated at 37°C and 5% CO2 for 1 hour. Plates were gently rocked by hand every 15 minutes. After incubation, 1 mL of 0.6% carboxymethylcellulose overlay was added to each well. Plates were incubated at 37°C and 5% CO_2_ for 4 days. On day 4, the overlay was aspirated. Wells were fixed with 10% neutral-buffered formalin and stained with 0.1% crystal violet before plaques were counted. Cell culture and plaque assay reagent recipes were adapted from Case, et al [26].

### SARS-CoV-2 challenge of K18-hACE2 mice

Stocks of the SARS-CoV-2 Delta variant B.1.617.2 hCoV-19/USA/WV- WVU-WV118685/2021 (GISAID Accession ID: EPI_ISL_1742834) were created from a patient sample at WVU that was propagated in Vero E6 cells (ATCC-CRL-1586) [27]. The stocks were sequenced to confirm there were no mutations. Omicron variant (strain BA.5) stocks were obtained from the labs of Dr. Luis Martinez-Sobrido and Dr. Jordi Torrelles at the Texas Biomedical Research Institute. At the time of challenge, vaccinated and control K18-hACE2 mice were anesthetized with an IP injection of ketamine/xylazine, then 25 µL of a 10^4^ PFU solution of Delta or 10^5^ PFU solution of Omicron virus was administered by pipette into each nare (50 µL total dose).

### Disease scoring of SARS-CoV-2 challenged mice

K18-hACE2 mice were evaluated every day after challenge to track disease progression through in-person health checks and using the SwifTAG video monitoring system. Rectal temperatures and weight measurements were recorded each day in addition to scores related to weight loss, changes in activity, appearance, eye closure/conjunctivitis, and respiration. Scores were awarded based on severity of disease phenotypes and follow a scale that has been described previously [17,25,27,28]. On each day, scores in each category were combined and recorded as one overall numerical score. Mice that received a total score of 5 or reached 20% weight loss before day 10 post-challenge were humanely euthanized.

### Mouse euthanasia and tissue collection

At day 2, 10, or due to meeting humane endpoint criteria, K18-hACE2 mice were euthanized with IP pentobarbitol (390mg/kg) (Patterson Veterinary 07-805-9296) followed by cardiac puncture. Blood from cardiac puncture was centrifuged to collect the serum for downstream analysis. Lung and brain tissue were dissected out for downstream histology, serology, and quantification of viral burden. Homogenization of lung and brain was performed following a previously established protocol [17,25]. A nasal wash was performed on each mouse by pushing PBS (1mL) by catheter through the nasal pharynx and collected for analysis. For RNA purification in BSL2, lung and brain homogenate as well as nasal wash was treated with TRIzol Reagent.

### SARS-CoV-2 plaque assay from tissue homogenate

Vero E6 ACE2/TMPRSS2 cells were plated at 150,000 cells per well in 12-well plates and incubated at 37°C and 5% CO2 for 24 hours. Mouse lungs collected at euthanasia were weighed then homogenized in 1mL PBS. Then, homogenate was centrifuged at 15,000 x g for 5 minutes. Supernatant from mouse lung homogenate was diluted in media 1:3, 1:10, then four ten-fold serial dilutions. Cell media was aspirated. 200μl of sample was added to each well, in duplicate. Plates were incubated at 37°C and 5% CO2 for 1 hour. Plates were gently rocked by hand every 15 minutes. After incubation, 2 mL of 0.6% carboxymethylcellulose overlay was added to each well. Plates were incubated at 37°C and 5% CO2 for 4 days. On day 4, the overlay was aspirated. Wells were fixed with 10% neutral-buffered formalin and stained with 0.1% crystal violet.

### qRT-PCR quantification of SARS-CoV-2 viral copy number in mouse tissues

RNA from nasal wash, lung, and brain homogenates of virus-challenged mice was purified using the the Direct-zol RNA miniprep kit (Zymo Research R2053) according to the manufacturer’s protocol. qPCR of the SARS-CoV-2 nucleocapsid gene was then performed for each mouse and sample using the Applied Biosystems TaqMan RNA to CT One Step Kit (ThermoFisher Scientific 4392938) to measure viral copy number via transcript number with specifications for each reaction that have been described previously [17,25,27,28].

### Measurement of CXCL10 concentrations in mouse serum and lung

Concentrations of CXCL10 in K18-hACE2 mice were measured in serum and lung supernatant collected at euthanasia using the Mouse Magnetic Luminex Assay kit for mouse CXCL10/IP-10/CRG2 (R&D Systems LXSAMSM-01). Both serum and supernatant samples were diluted 1:2 in the kit’s Calibrator Diluent RD6-52 then utilized in the assay procedure provided by the manufacturer. The assay plate was read on the Luminex MAGPIX instrument and chemokine levels were quantified based on a standard curve.

### Histopathological evaluation of lung tissue inflammation

The left lobes of mouse lungs were collected at euthanasia and stored in 10% neutral buffered formalin for one week to fix. Fixed tissues were sectioned and mounted on slides, then Hematoxylin and Eosin stained for analysis. Primary histopathological scoring was performed by iHisto under the supervision of chief pathologist Michelle X. Yang, MD, PhD. H&E-stained slides were evaluated for acute and chronic inflammation. Acute inflammation was marked by the infiltration of neutrophils in the parenchyma, blood vessels, and airways. Chronic inflammation was marked by mononuclear infiltrates in the same areas of the tissue. Semiquantitative scores for each condition (0- none, 1- minimal, 2- mild, 3- moderate, 4- marked, 5- severe) were awarded for tissue from each mouse. Additional analysis was performed by West Virginia University’s Department of Pathology to identify more finite characteristics of inflammatory pathology. Margination of inflammatory cells in blood vessels was evaluated and graded as none, mild, moderate or severe. Viral cytopathic changes in the epithelial and interstitium were also evaluated and graded as absent, mild, moderate or severe. Other parameters that were evaluated included type of inflammatory cells, distribution of the inflammatory aggregates, presence of pneumonic infiltrates, bronchiolitis and vasculitis.

### Statistical analyses

The statistical analysis of data sets in this study was performed using GraphPad Prism version 9. Mouse experiments were performed with an n=10 for all groups, with n=3 mice per group euthanized on day 2 and n=7 mice euthanized at day 10 post-challenge. Normally distributed data sets were compared using ordinary one-way ANOVA with Tukey’s multiple comparisons tests. Kaplan-Meier survival curves were created to compare the survival of vaccinated K18- hACE2 mice following viral challenge. Unpaired t-tests were utilized to compare two data sets.

## Results

### VLP-RBD adjuvanted with BECC 470 (TLR4 agonist) induces IgG and IgA antibody production when administered via the nasal route

SARS-CoV-2 infection induces mucosal immune responses including production of IgA antibodies. To assess mucosal responses as well as systemic antibodies elicited by our novel VLP-RBD-BECC vaccine, K18-hACE2 transgenic mice were primed via the intranasal (IN) route using 10 µg VLP-RBD formulated with 25 µg BECC 470 (Fig. 1A). A control group of mice were vaccinated via the intramuscular (IM) route with 1/10^th^ of a human dose (10 μg) of mRNA-1273-- a highly protective Spike protein-based COVID-19 vaccine that has been used to vaccinate a large portion of the United States population. After four weeks, mice were administered a second identical boost dose of either vaccine to mimic the human COVID-19 vaccine schedule. We measured levels of IgG specific to the SARS-CoV-2 Spike protein receptor binding domain (RBD) at four weeks post-prime (Fig.1B), four weeks post-boost (Fig. 1C), and after challenge with the SARS-CoV-2 Delta strain (Fig. 1D). IM mRNA-1273 vaccinated mice showed high IgG levels post-prime whereas IN VLP-RBD-BECC mice showed relatively low early antibody responses. However, at post-boost the antibody levels in IN mice rose to levels near IM mRNA, albeit still significantly lower (Fig. 1C). At 10 days post-challenge with Delta, RBD specific IgG levels from IN VLP-RBD-BECC or mRNA remained similar to post-boost (no significance) (Fig. 1D). mRNA vaccines are known to induce low levels of IgA in serum [29,30]; however, it is not clear if mRNA induces mucosal antibodies in mice. At 10 days post-challenge, low RBD-specific IgA levels were indeed detectable in the serum of IM mRNA vaccinated mice (Fig. 1E). vaccination with IN VLP-RBD-BECC, however, induced anti-RBD IgA antibodies at a much greater level in serum (Fig.1E), nasal wash (Fig. 1F) and lung supernatants (Fig. 1G) post-challenge. These data confirm that intranasal vaccination with VLP-RBD-BECC can induce RBD-specific IgG antibodies in addition to highly desirable IgA antibodies.

**Figure 1:**
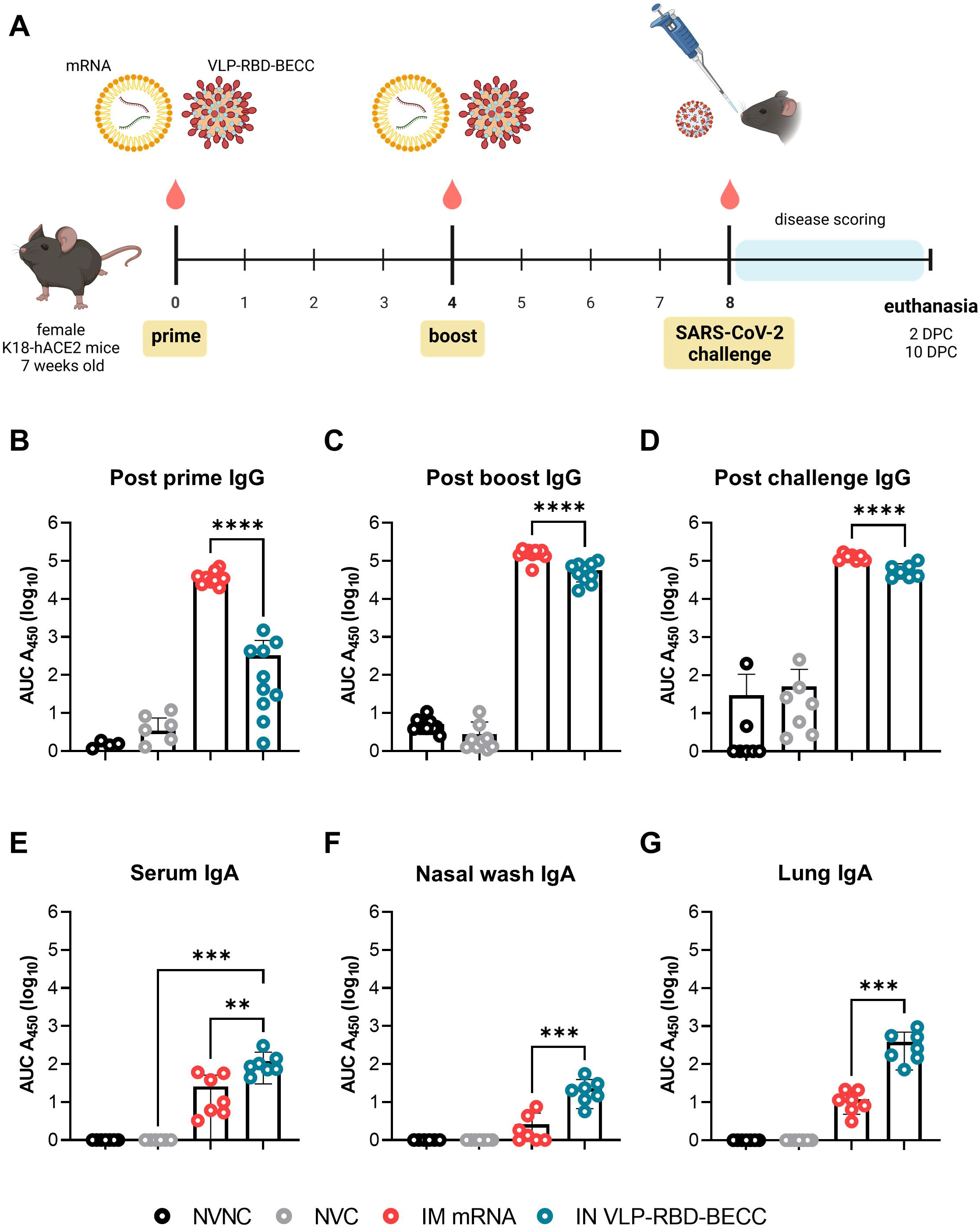
Intranasal VLP-RBD-BECC induces strong antibody responses in K18-hACE2 mice. Anti-RBD IgG levels were measured in serum A) four weeks post-prime vaccination, and B) four weeks post-boost vaccination (n=10 mice per group). Serum, nasal wash, and lung supernatant were collected at euthanasia points (NVC mice reached total morbidity by day 6 while all other groups survived to day 10 post-challenge) to measure C) serum anti-RBD IgG, D) serum anti-RBD IgA, E) nasal wash anti-RBD IgA, and F) lung supernatant anti-RBD IgA (n=7 mice euthanized at 6 or 10 DPC). Each point denotes one biological replicate. One Way ANOVA with Tukey’s Multiple Comparisons was performed to determine *P* value: \*\*\*\**P*<0.0001, \*\*\**P<0.0006, **P*=0.0063

### IN VLP-RBD-BECC vaccinated K18-hACE2 mice produce broadly neutralizing antibodies against SARS-CoV-2 variants

One tenet of highly effective antibody responses is pathogen neutralization which blocks host cell receptor binding for viral entry. Antibodies raised in response to natural infection or vaccination bind the SARS-CoV-2 spike protein to block host cell ACE2 receptor binding and thus inhibit downstream viral replication and the progression of infection. To assess whether or not anti-RBD IgG antibodies from IN VLP-RBD-BECC could execute this inhibition, we performed an *in vitro* ACE2-RBD binding inhibition assay using the serum from vaccinated K18- hACE2 at euthanasia post-challenge. Serum IgG antibodies from IM mRNA vaccinated mice bound RBD from the ancestral, Alpha, Beta, and Delta SARS-CoV-2 variants of concern (VOC) and inhibited ACE2 binding at a greater capacity than serum from IN VLP-RBD-BECC mice *in vitro* (Fig. 2A). VLP-RBD-BECC antibodies at a 1:500 dilution still inhibited greater than 50% ACE2 binding to all VOC (Fig. 2A). At the greatest dilution, 1:5000, serum from both vaccinated groups dropped below 50% binding inhibition. It should be noted that mRNA immunized mice have the highest concentration of RBD-binding antibodies in serum post-vaccination and challenge (Fig. 1). IgG antibodies from VLP-RBD-BECC showed a calculated percent binding PRNT50 of 2341 against Delta RBD which was not dramatically reduced compared to the PRNT50 of mRNA-elicited antibodies at 3741 (Fig.2B). Authentic virus neutralization was further evaluated in a viral plaque reduction assay where the SARS-CoV-2 Delta variant was propagated *in vitro* with decreasing concentrations of serum from vaccinated mice collected post-boost. At a low dilution (1:10), mRNA vaccinated mouse serum fully prevented plaque formation (100% reduction) and at higher dilutions (1:100 and 1:1000) still reduced plaque formation by 88% or more (Fig. 3). Serum from VLP-RBD-BECC vaccinated mice was also highly effective at neutralizing virus to prevent plaque formation when added to culture media at a 1:10 dilution and reduced more than 88% or 78% of plaques at higher dilutions (1:100 and 1:1000 respectively). These assays show that although IgG antibodies elicited by our VLP-RBD- BECC vaccine have reduced inhibitory activity compared to those from mRNA, total antibodies (which may include IgM, IgA and other antibody subclasses) from these mice are still highly effective at limiting SARS-CoV-2 viral replication and plaque formation *in vitro*.

**Figure 2:**
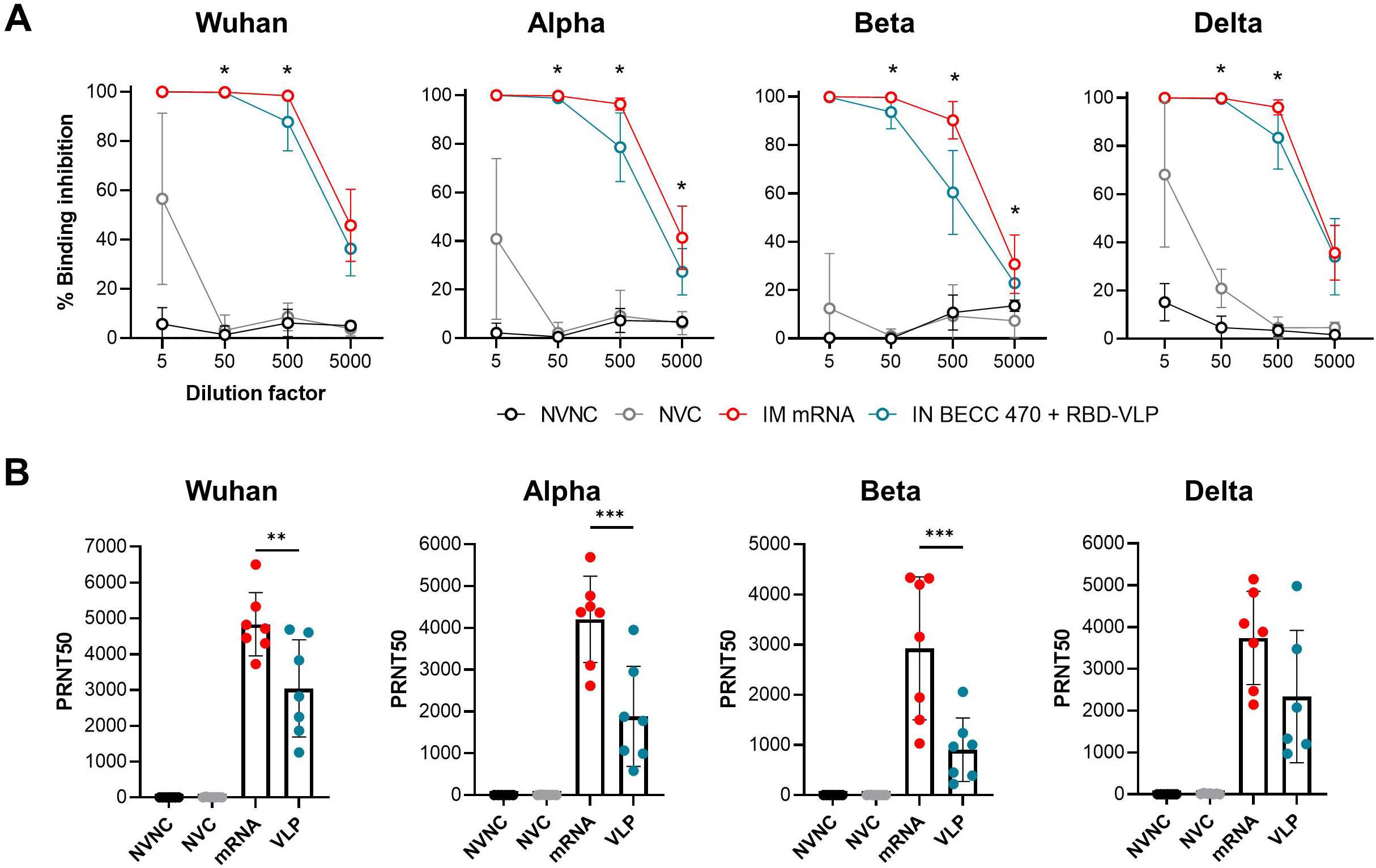
Neutralizing antibodies against multiple SARS-CoV-2 VOC RBD are elicited from IN VLP-RBD-BECC. A) Serum IgG antibodies from mRNA or VLP-RBD-BECC vaccinated mice were analyzed at euthanasia point 6 (NVC) or 10 (IM mRNA or IN VLP-RBD-BECC) days post-challenge to measure *in vitro* ACE2-RBD % binding inhibition and compared to PBS- vaccinated and challenged mice. Points represent the average of n=6-7 biological replicates. Unpaired T tests were performed for statistical analysis: * = significance between mRNA and VLP-RBD-BECC at dilution point (*P*<0.05). B) The PRNT50 (percent neutralization 50) values were interpolated for each biological replicate from the dilution curve of % binding inhibition. way ANOVA with Tukey’s Multiple Comparisons was performed for statistical analysis. \*\*\**P*<0.0091, \*\**P*=0.0020.

**Figure 3:**
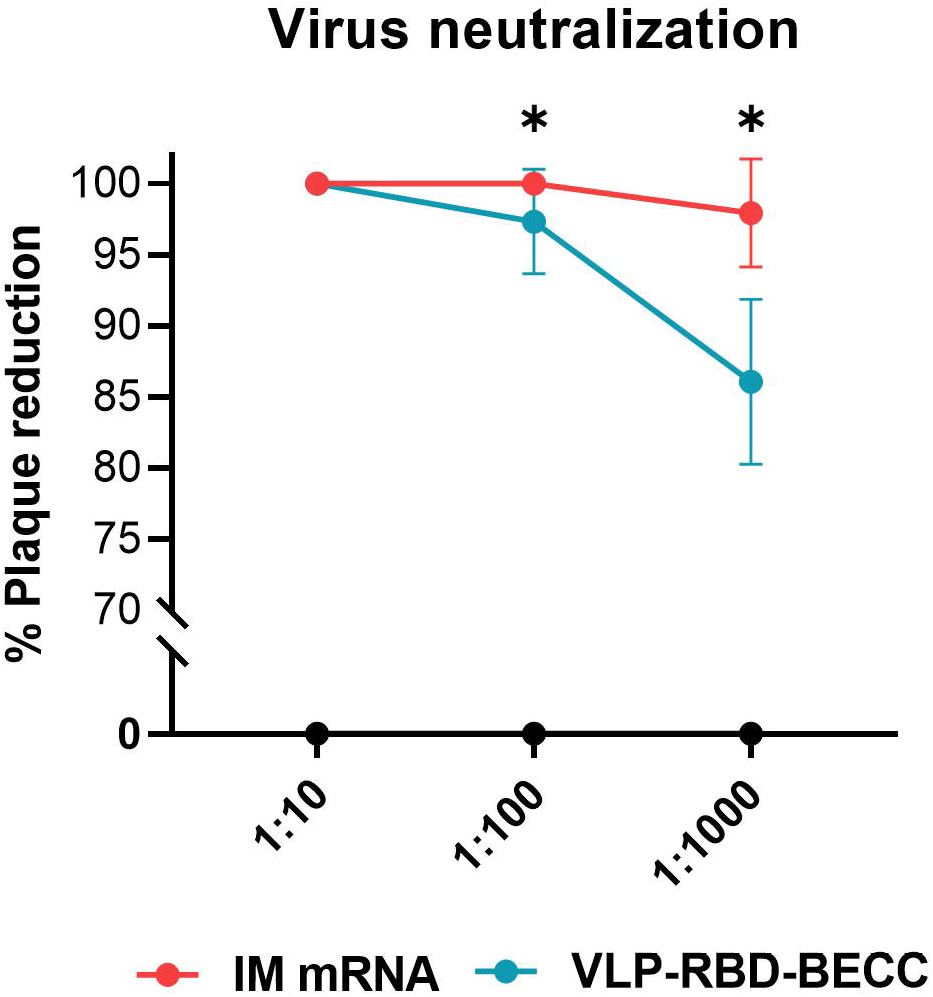
**Serum from VLP-RBD-BECC vaccinated mice reduce plaque formation i*n vitro.*** Serum collected 4 weeks post-boost was added with Delta SARS-CoV-2 (n=10 biological samples per group) to Vero E6-AT cells to measure the ability of antibodies to reduce plaque formation. Unpaired t-tests were performed for statistical analysis: * *P*<0.05

### IN VLP-RBD-BECC protects mice against challenge with SARS-CoV-2 Delta variant

Each major SARS-CoV-2 VOC to emerge after the ancestral Wuhan strain, has presented pathological differences in the K18-hACE2 challenge model [27,31–33]. In particular, we have observed that the Delta variant requires a slightly higher dose to cause 100% mortality, but also that the tissue inflammation signatures and immunological response in mice are significantly different than other variants [27]. We challenged the IN VLP-RBD-BECC and IM mRNA-1273 vaccinated K18-hACE2 mice to evaluate protection conferred against the Delta variant. Four weeks after boosting, mice were intranasally challenged them with a lethal dose (10^4^ PFU) of the SARS-COV-2 Delta variant to measure protection (Fig. 1A). Intranasal and intramuscular vaccines were equally matched in their disease-limiting abilities (Fig. 4). Compared to PBS-vaccinated and challenged mice, IM mRNA and IN VLP-RBD-BECC vaccinated mice maintained low disease scores over the course of the 10-day challenge window (Fig. 4A). IN VLP-RBD-BECC mice did not experience dramatic weight loss or drops in temperature, similar to the IM mRNA group (Fig. 4BC). VLP-RBD-BECC also conferred 100% survival compared to non-vaccinated mice which reached total morbidity and required humane euthanasia by day 6 post-challenge (Fig. 4D). This data together suggests that VLP-RBD-BECC administered by two intranasal doses is as effective as mRNA-1273 at conferring protection against morbidity and mortality in SARS-CoV-2 related disease in K18-hACE2-mice.

**Figure 4:**
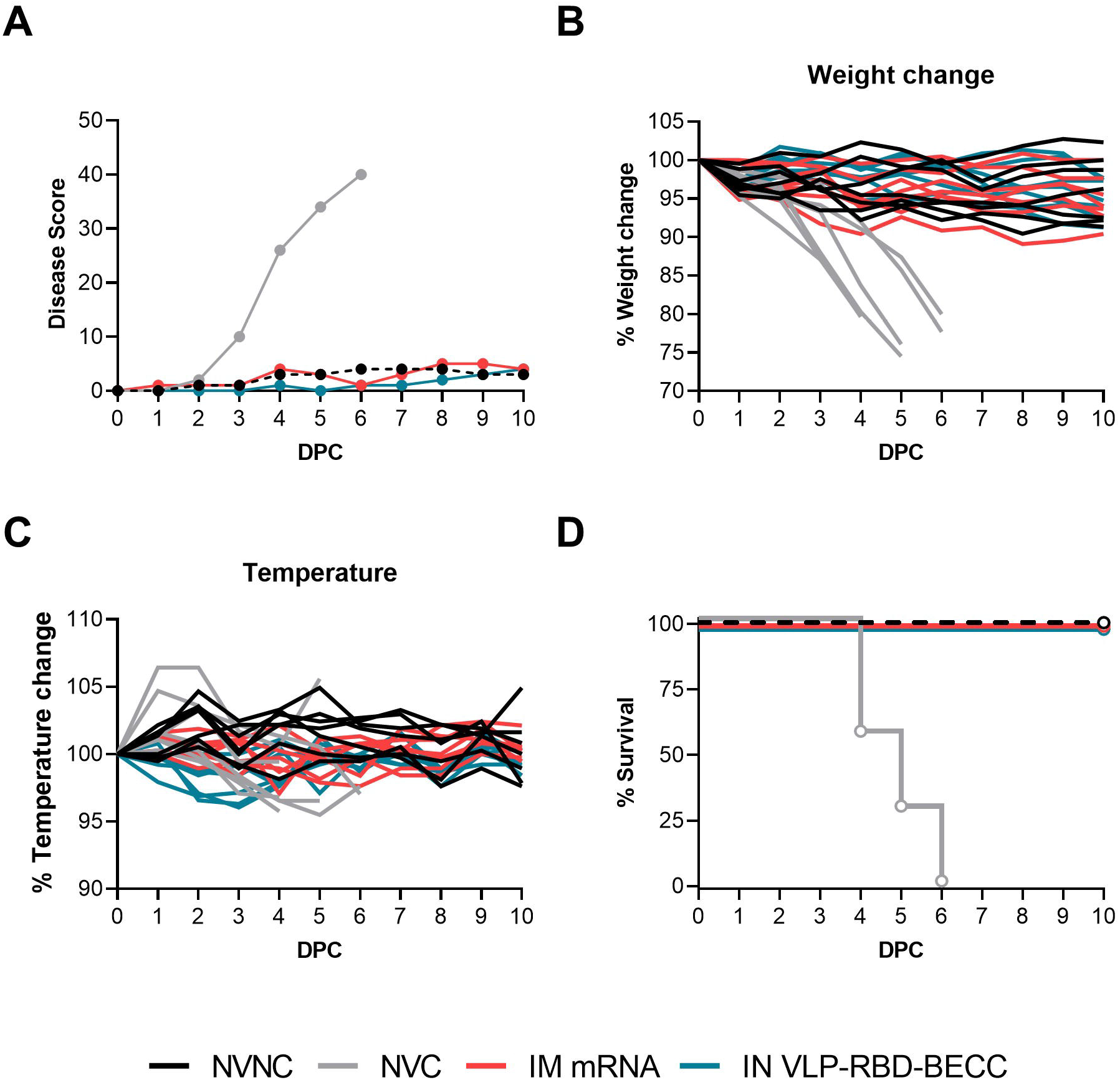
Vaccination with VLP-RBD-BECC intranasally confers protection against lethal challenge with Delta. A) Mice were monitored daily for disease progression after challenge with the sum of their disease scores reported daily for up to 10 days post-challenge. B) Daily weight and C) temperature change was measured to monitor disease. Mice in the NVC group reached morbidity by day 6 at which point disease scores stopped being reported. Any mice euthanized before complete euthanasia of the group had their score retained for reporting. D) Kaplan Meier survival curve of vaccinated and viral-challenged K18-hACE2 mice. (n=7 mice per group)

### SARS-CoV-2 viral replication is limited by intranasal VLP-RBD-BECC

In our challenge study, VLP-RBD-BECC and mRNA vaccinated K18-hACE2 mice were euthanized at day 2 and day 10 post-Delta challenge to assess the vaccines’ ability to limit viral replication. Authentic plaque assays using lung supernatant from the mice euthanized at day 2 showed that IN VLP-RBD-BECC vaccination and IM mRNA significantly limited viral replication in the lung compared to the PBS-vaccination control (Fig. 5A). Additional qRT-PCR analyses of the mice’s nasal wash, lung and brain tissues further supported this finding. SARS-CoV-2 viral nucleocapsid RNA copies were significantly lower in the nasal wash of VLP-RBD-BECC mice compared to PBS vaccinated mice at day 2 and was further reduced at day 10 (Fig. 5B). In the lung, viral RNA burden was significantly lower in mRNA vaccinated mice than the PBS group, however there was no significant reduction in the viral RNA burden of VLP-RBD-BECC lungs at day 2 (Fig. 5C). At day 10, viral RNA burden in the lungs in both vaccine groups had reduced significantly from the level of controls, although it remained slightly above the limit of detection (Fig. 5C). Both vaccines prevented dissemination to and detection of viral RNA in the brain at 10 days post-challenge when it was detectable in the brain tissue of control mice (Fig. 5D).

**Figure 5:**
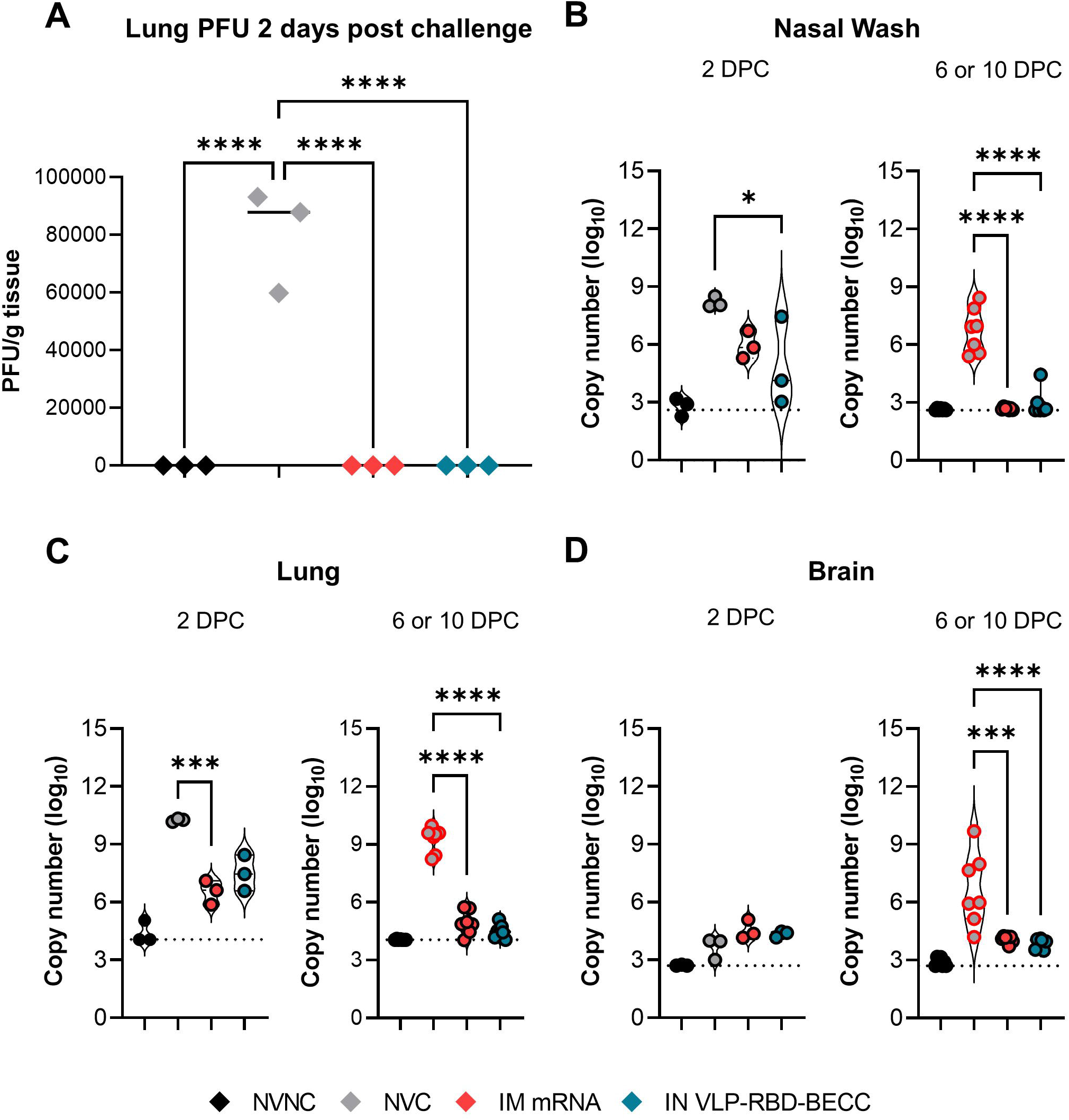
IN VLP-RBD-BECC and IM mRNA limit viral replication and tissue viral burden. A) PFUs from lung homogenates of mice euthanized 2 days post-challenge (n=3 mice per group). B) qRT-PCR was performed to measure viral RNAs in the nasal wash, C) lung and D) brain of mice in each experimental group euthanized at day 2 post-challenge or at euthanasia. Mice in the NVC group all reached morbidity by day 6 (symbols outlined in red) while all other groups survived to day 10. Copy numbers calculated below the limit of detection (LOD designated by dashed line) were set as equal to the LOD for reporting. One way ANOVA with Tukey’s Multiple Comparisons was performed for statistical analysis. \*\*\*\**P*<0.0001, \*\*\**P*=0.0003, \*\**P*=0.0066, *\*P*=0.0445. (n=3 mice per group euthanized at 2 DPC, n=7 mice euthanized humane endpoint or 10 DPC)

### Intranasal vaccination limits the production of CXCL10 following SARS-CoV-2 challenge in K18-hACE2 mice

CXCL10 is a potent mediator of inflammation and immune cell homing to the tissues during SARS-CoV-2 infection [34]. It’s production in the lung is linked to the cytokine storm experienced by hospitalized patients with severe cases of COVID-19 [35]. Previously, we identified that K18-hACE2 mice have high concentrations of CXCL10 in the lung 6 days post delta variant challenge [27]. To assess if vaccination strategies to protect individuals against severe COVID-19, including VLP-RBD-BECC, confer protection in part by limiting the production of the chemokine, we quantified CXCL10 levels in the serum and lung supernatant of vaccinated mice after Delta SARS-CoV-2 challenge. In the serum, CXCL10 production increased at day 2 after viral challenge in PBS vaccinated control mice (Fig.6A). By day 10, this concentration had reduced slightly but at both timepoints, the concentration of the chemokine in IM mRNA or IN VLP-RBD-BECC remained low. CXCL10 production also peaked in the lung tissue of PBS vaccinated mice at day 2, to much higher concentrations than were detected in the serum (Fig.6B). In vaccinated mice, CXCL10 concentrations in the lung were also limited with no significant change between timepoints. The lack of CXCL10 production seen in the serum and lungs of vaccinated mice when compared to controls, demonstrate that vaccination effectively limits the production of proinflammatory chemokines produced in response to viral challenge.

**Figure 6:**
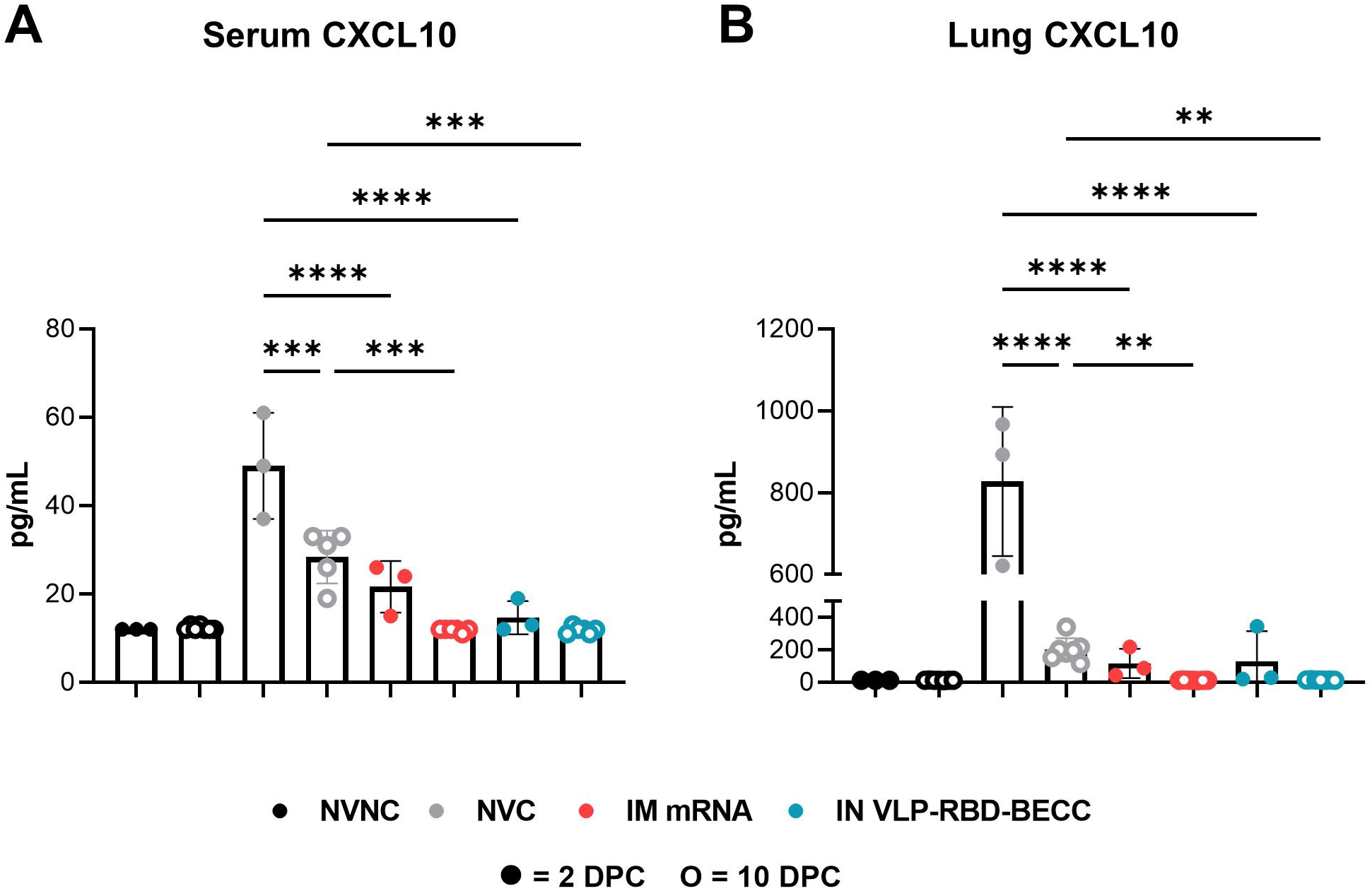
Vaccination limits CXCL10 production in the lung after Delta variant challenge. CXCL10 concentrations were quantified in the A) serum and B) lung supernatant of K18-hACE2 mice 2 or 10 days after SARS-CoV-2 viral challenge (NVC mice surviving past 2 DPC were euthanized 4-6 DPC). Each symbol represents one biological replicate. One way ANOVA with Tukey’s Multiple Comparisons was performed for statistical analysis. \*\*\*\**P*<0.0001, \*\*\**P<*0.0003 \*\**P*<0.0013

### Intranasal VLP-RBD-BECC vaccination ameliorates lung inflammation in K18-hACE2 mice after Delta challenge

As a final aspect of vaccine evaluation, lung inflammation was measured by histopathological scoring of acute and chronic inflammation phenotypes. No significant inflammation was seen in the lungs of mice in the control group which served as a comparison to vaccine groups (Fig. 67A). Delta challenge in PBS-vaccinated mice lead to viral cytopathic and reactive-proliferative changes in the epithelia of the terminal bronchioles, alveoli, and interstitial cells. Marked margination of inflammatory cells in blood vessels was observed and associated with a diffuse lymphocyte and histiocyte rich inflammatory infiltrate that involved 50-75% of the pulmonary parenchyma. At 2 days post-challenge, intramuscularly vaccinated mice showed higher acute inflammation scores than the intranasal group, suggesting that our intranasal vaccine better protects mice from early development of lung infection than IM mRNA, perhaps due to the route of administration and localized immune responses (Fig.7B). Interestingly, PBS vaccinated mice showed much lower lung inflammation at this time point than IM mRNA mice (there was no significant difference between PBS and IN VLP-RBD-BECC), suggesting perhaps a delayed inflammatory response to virus in the tissue. IM mRNA vaccinated mice euthanized 10 days post-challenge were given low scores for acute inflammation which were not statistically different from IN VLP-RBD-BECC mice (Fig.7C). The diminished extent of pulmonary involvement was further supported by the observation of no significant margination of inflammatory cells or viral cytopathic changes in the vaccinated groups which decreased by day 10 (Fig.7D). Overall, mice that were vaccinated either with IM mRNA or IN VLP-RBD-BECC showed marked attenuation of total inflammation compared to PBS vaccination, and involved less than 25% of the parenchyma. Inflammation in vaccinated mice consisted predominantly of tight, peribronchial and perivascular lymphocyte-rich aggregates (Fig.7EF). In total this histopathological analysis shows that IN VLP-RBD-BECC is able to control lung inflammation in the lungs of mice throughout the post-challenge window against SARS-CoV-2 related lung inflammation.

**Figure 7:**
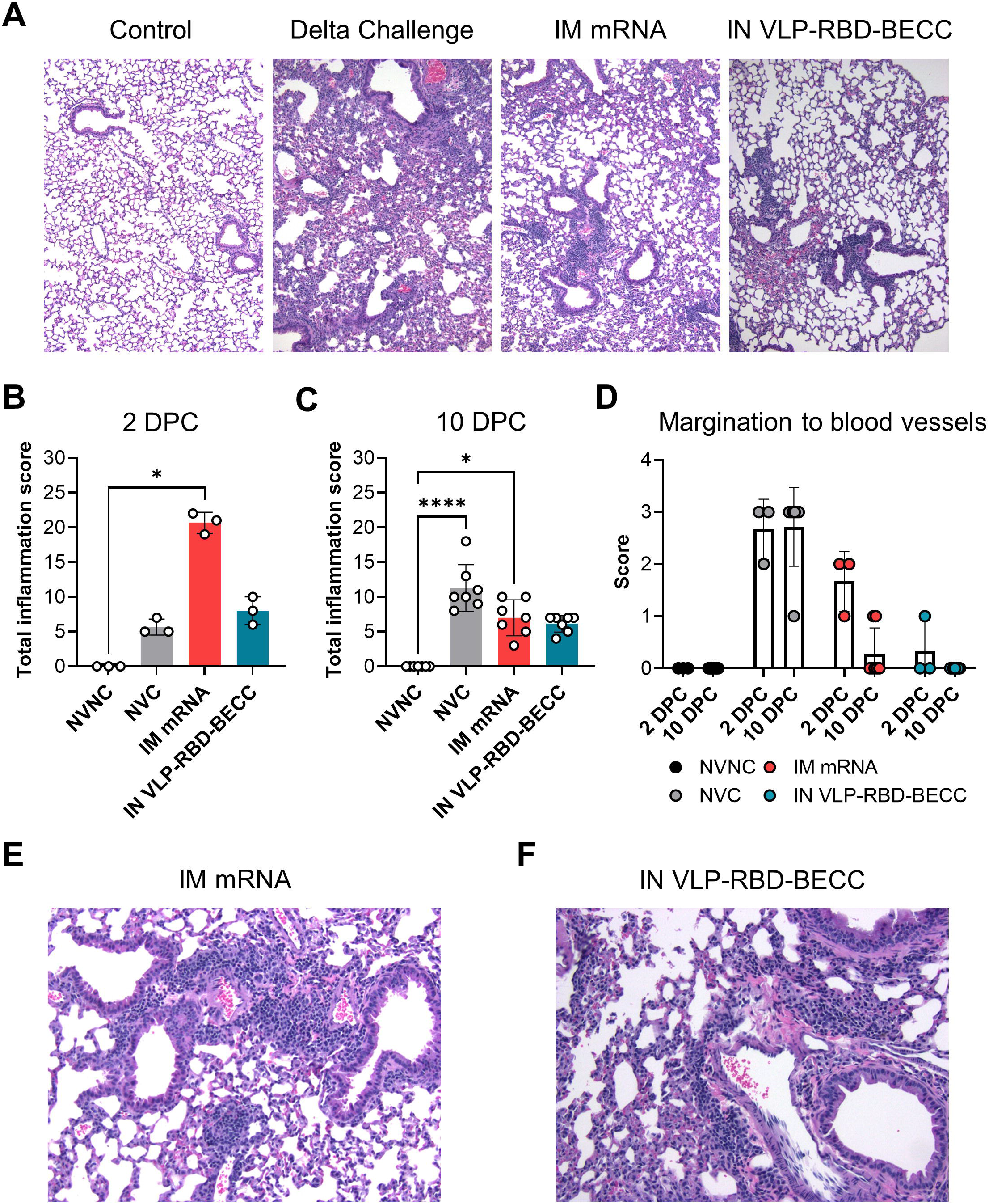
Histopathological analysis shows lung inflammation is limited by VLP-RBD- BECC in SARS-CoV-2 challenged mice. A) Representative images of H&E-stained mouse lungs collected at euthanasia after viral challenge. Total inflammation scores comprised of acute and chronic inflammation scores were awarded to mice euthanized B) two (n=3) or C) 10 days (n=7) post-challenge (NVC mice surviving past 2 DPC were euthanized on or before 6 DPC). For each parameter of acute and chronic inflammation, 0=none; 1=minimal; 2=mild; 3=moderate; 4=marked; 5=severe. D) Margination of immune cells into the blood vessels was scored for each mouse where 0=none; 1=mild; 2=moderate; 3=severe. Statistical analyses were performed using one way ANOVA with Tukey’s Multiple Comparisons: \*\*\*\**P*<0.0001, \**P*<0.0295

### Mice are less protected by intranasal VLP-RBD-BECC against Omicron challenge

The rise of the SARS-CoV-2 Omicron variant to clinical dominance resulted in a reduction of the protection that vaccines formulated against ancestral strains of the virus could provide. Using serum collected at euthanasia from IN VLP-RBD-BECC mice challenged with the Delta variant, we measured the capacity of IgG antibodies to bind Omicron RBD and thus prevent ACE2 binding. VLP-RBD-BECC -elicited antibodies showed reduced RBD binding against the variant at dilutions of 1:5 and 1:50 compared to antibodies from mRNA-vaccinated mice (Fig.8A). To determine how this reduced neutralization activity correlated to a potential reduction in protection against the SARS-CoV-2 Omicron virus, we vaccinated a new cohort of mice with two doses of mRNA intramuscularly, or VLP-RBD-BECC intranasally. After prime, VLP-RBD-BECC mice again showed lower amounts of anti-Wuhan RBD IgG antibodies in serum, which increased post-boost (Fig.8B). The utility of matching vaccines to viral variant has been argued as a superior method of providing protection, resulting in the formulation of bivalent (multi-variant) mRNA vaccines for COVID-19 [36,37]. We repeated antibody quantification ELISAs using RBD from the Omicron variant and observed a reduction in anti-RBD IgG antibodies in mRNA-vaccinated serum post-prime, and VLP-RBD-BECC serum at both time points (Fig.8C). Omicron does not cause morbidity in the K18-hACE2 transgenic mouse model, negating survival as a means of measuring protection in vaccine studies. To assess protection from VLP-RBD-BECC compared to mRNA, mice were euthanized at 2 and 6 days after intranasal challenge with 10^5^ PFU Omicron to evaluate a reduction in viral replication. qRT-PCR analysis of the viral nucleocapsid gene showed no difference between control and vaccinated groups in the nasal wash or brain (data not shown). In the lung, viral RNA copy numbers were not significantly different between Omicron-challenged control mice and IN VLP-RBD-BECC groups at day 2 or day 6 (Fig.8D). However, viral replication, measured by plaque forming units in mouse lung tissue, was significantly lower at day 2 in VLP-RBD-BECC mice than in the lung tissue of controls (Fig.8E). By day 6, actively replicating virus was undetectable by plaque assay in any group. These data together demonstrate that although our VLP-RBD-BECC vaccine formulated using RBD from an ancestral virus elicits an antibody response that less-efficiently targets the Omicron variant, the vaccine still reduces the development of viral burden in the lung tissue of mice.

**Figure 8:**
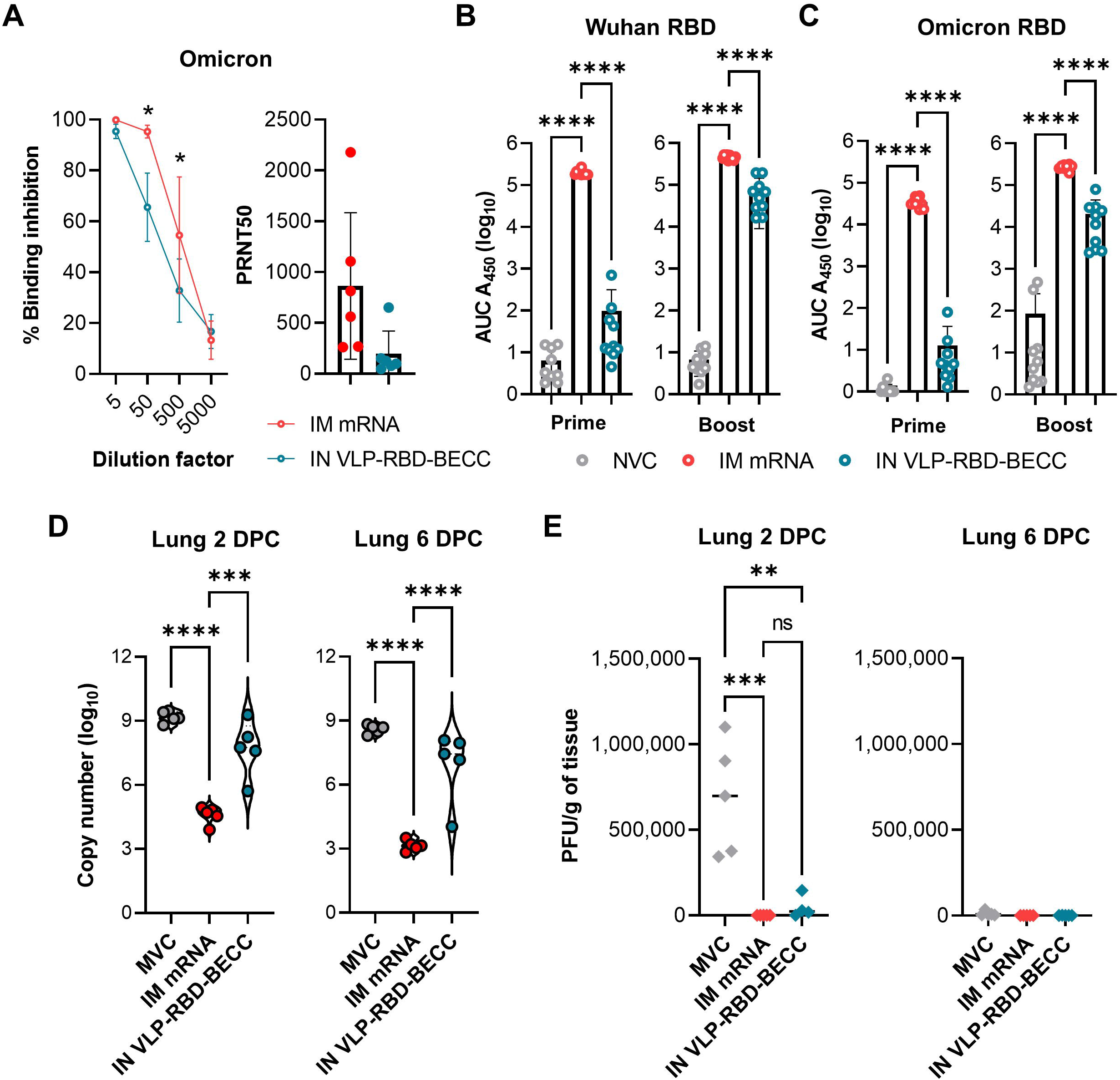
Protection from VLP-RBD-BECC is reduced against Omicron challenge. A) *In vitro* binding inhibition assays were performed to measure the binding of serum IgG antibodies collected two weeks post boost to Omicron RBD. Points denote the average of n=6 biological replicates, Unpaired T tests were performed for statistical analysis: * = significance between mRNA and VLP-RBD-BECC at dilution point (*P*<0.05). B) Anti-Wuhan RBD IgG antibodies and C) anti-Omicron RBD IgG antibodies were quantified by ELISA in serum two weeks post prime and two weeks post boost. D) qRT-PCR analysis was performed to determine copy number of the SARS-CoV-2 nucleocapsid gene in lung tissue homogenates collected 2- or 6-days post-challenge. Dashed lines indicate the limit of detection. E) Plaque assays were performed using lung tissue supernatant collected 2- or 6-days post-challenge to measure viral burden. Each point denotes one biological replicate. One Way ANOVA with Tukey’s Multiple Comparisons was performed to determine *P* value: \*\*\*\**P*<0.0001, \*\*\**P*<0.0006, *\*\*P*=0.0017

## Discussion

As of the Fall of 2022, all COVID-19 vaccines approved for human use in the United States are administered via the intramuscular route. Across the globe, however, countries such as China and India have begun to approve a small number of intranasally-delivered platforms for use in the continued fight against COVID-19, and even more are being actively evaluated in preclinical and clinical trials [11]. Bharat Biotech’s adenovirus vectored vaccine (formerly ChAd-SARS-CoV-2-S, now BBV154) after extensive preclinical characterization was approved in India for both primary vaccination and as a boost [38–40]. Another adenovirus vectored vaccine (Ad5- nCoV-S) by CanSino Biologics in China was approved for use as an inhalable booster after initial approval for intramuscular use, highlighting the potential to explore alternative routes of administration with current vaccines to successfully incorporate intranasal vaccines into existing vaccine schedules [41,42]. While administration of mRNA intranasally would be an attractive approach to implementing an intranasal vaccine by repurposing an approved vaccine formulation, the lipid nanoparticle shells of these vaccines are not formulated for immunogenicity in the mucosa [12]. For intranasal use, vaccine platforms based on virus-like particles which are highly immunogenic and incapable of replication (unlike adenoviruses or lentiviruses making them safe for the elderly or immunocompromised), are a promising future technology [43]. SpyBiotech developed a VLP-based vaccine for COVID-19 that, when adjuvanted with alum or alum+CpG, conferred protection to rhesus macaque following SARS-CoV-2 challenge [23,44]. The preclinical efficacy of this vaccine quickly prompted its advancement to clinical trials, and here we have evaluated this antigen by intranasal route and in combination with the BECC adjuvant.

In our study, we evaluated the combinatorial use of SpyBiotech’s VLP-RBD, and the BECC 470 adjuvant, established by our lab to be an effective intranasal vaccine antigen in BReC-CoV-2, as an intranasal vaccine for COVID-19. K18-hACE2 mice vaccinated with two doses of intranasal VLP-RBD-BECC were protected against the development of severe disease and morbidity after challenge with a lethal dose of the SARS-CoV-2 Delta variant. We demonstrated that intranasal VLP-RBD-BECC elicits high anti-RBD IgG and IgA production, as compared to intramuscular mRNA. The neutralizing activity of serum antibodies was lower in IN mice than IM, but IN vaccination was still able to limit viral replication at 2 DPC. Viral RNA burden in the lungs and nasal wash was reduced by IN vaccination at 10 DPC as compared to in NVC mice. Importantly, IN vaccination prevented the dissemination of virus into the brain 10 DPC. Lung inflammation, conferred potentially by the reduced production of the chemokine CXCL10, was consistently limited after IN vaccination over the course of challenge studies (2 and 10 DPC).

Despite similarities in the viral RNA burden detectable in the lungs of IN and IM vaccinated mice and similar scores for inflammation at day 10 post-challenge, IN VLP-RBD-BECC mice displayed lower scores for lung inflammation at day 2 when compared to IM mRNA. Together our data demonstrates that VLP-RBD-BECC administered via the intranasal route can provide similar protection to mRNA-1273 against the Delta variant when administered intramuscularly. While both intramuscular mRNA and intranasal VLP-RBD-BECC were able to fully protect mice from challenge, future studies should evaluate the dose ranges and longevity of protection for each vaccine formulation and vaccination scheme.

The hepatitis B surface antigen-based VLP-RBD used in this vaccine study utilized RBD from an ancestral strain of SARS-CoV-2 (Wuhan/Washington-1). While currently approved COVID-19 vaccines on the market have utilized ancestral sequences and proteins of RBD and Spike, the concept of matching vaccine antigen to challenge strain has recently gained more attention. Emerging variants of SARS-CoV-2 acquire additional mutations to the viral Spike protein which decreases the ability of neutralizing antibodies and other aspects of the immune system that recognize the ancestral protein to recognize new variants like Omicron. Moderna and Pfizer-BioNTech’s bivalent mRNA booster vaccines were recently approved for use in the United States as a means of enhancing variant-specific immunity [36,37]. Both vaccines conserve inclusion of genetic material from the ancestral strain, but additionally contain genetic sequences encoding the Omicron BA.4 and BA.5 variants. It’s been largely reported that original COVID-19 vaccines remained partially protective against severe disease against the Delta and Omicron variants albeit with increased waning of protection over time [45]. We showed previously that VLP-RBD with other adjuvants was protective intramuscularly against early SARS-CoV-2 variants of concern [17]. Intranasal VLP-RBD-BECC was 100% protective against lethal Delta challenge in the K18-hACE2 mouse, boasting the superiority of this vaccine as compared to our BreC-CoV2 vaccine [16]. One area where intranasal VLP-RBD-BECC did not outperform intramuscular mRNA was in *in vitro* RBD-ACE2 neutralization. Although neutralizing antibodies from intranasal mice were high in serum post-boost and prevented Delta plaque formation, and our *in vitro* binding assay showed high ACE2-RBD binding inhibition against Delta when it was the challenge strain, vaccination-induced antibodies did not highly recognize Omicron RBD and did not convey sterilizing immunity against early Omicron virus replication in the lungs (at day 2). To meet our goal of developing a vaccination scheme that will improve protection against highly transmissible viral variants, future studies with this platform would need to assess the incorporation of VOC specific RBD into the VLP antigen before the vaccine moves into other preclinical models like hamsters where we can assess additional correlates of protection like reduced transmission.

Numerous intranasal vaccines utilizing a variety of platforms have been characterized in preclinical K18-hACE2 mouse challenge experiments [14,16,54–58,46–53]. Intranasal vaccines are also being developed by other labs to improve COVID-19 vaccines, although none reported so far have utilized the virus-like particle, and none have been evaluated preclinically against the highly virulent and pathogenic Delta variant. The adenovirus platform is another popular option for intranasal vaccines due to its high immunogenicity. One dose of ChAd-SARS-CoV-2- S, an adenovirus-based Spike protein-encoding vaccine that has progressed to evaluation in human trials, was shown to be protective in mice, hamsters, and rhesus macaques [39,40,57]. Hassan, et al. reported that ChAd-SARS-CoV-2-S confers lasting immunity against SARS-CoV- 2 variants of concern using serological assays against variant strain RBD. Although dampened compared to ancestral strains, ChAd-SARS-CoV-2-S induced neutralizing antibodies that bound Delta [57]. To our knowledge, this vaccine and its correlates of protection have yet to be evaluated authentically in a challenge study with the Delta variant. Vesin et al. evaluated the protection of a lentiviral based Spike vaccine (Lv:S) as a boost in Ad5:hACE2 transfected mice later challenged with the Delta variant [56]. Although this Delta challenge model did not utilize the K18-hACE2 transgenic mouse that is susceptible to SARS-CoV-2 induced pathology, it did show effective limitation of viral RNA post-challenge in the lung [56]. In these studies, researchers assessed the added benefits of intranasal vaccination that exist in addition to protection against morbidity and mortality. Immunogenicity studies for intranasal vaccines have described that they are able to not only elicit high mucosal antibody levels, but important T cell populations resident to the lung and respiratory tract [59]. This priming of the mucosa is likely a major contributor to reducing early disease pathology—reduced viral replication in the upper respiratory tract, limited lung inflammation, and decreased transmission/viral shedding—seen by our lab and others. Vesin et al. evaluated the implementation of their IN Lv:S vaccine as a booster in addition to a previously administered mRNA schedule [56]. This is an important variable to consider as we work to implement intranasal vaccines into the long-term COVID-19 response as most of the world will be intramuscularly vaccinated and require additional boosters. VLP-RBD-BECC could be an advantageous booster for COVID-19 protection due its mucosal IgA responses which would supplement the strong IgG-dominant responses incurred by mRNA priming. While VLP-RBD-BECC and mRNA were equally matched in their ability to confer protection against SARS-CoV-2 challenge, we showed that antibody responses to both vaccines were distinct. VLP-RBD-BECC as an intranasal vaccine primed the host for IgA production, which was not seen in mRNA vaccinated mice, but mRNA vaccinated mice showed higher IgG production systemically. IgA responses may provide better sterilizing immunity in the mucosa, protecting against disease pathology and possibly even transmission in a manner which systemic IgG levels cannot. The function of VLP-RBD-BECC induced IgA compared to mRNA-induced IgG merits further study. Additionally, the BECC 470 adjuvant used in this formulation was described to confer a Th1 dominant T cell response in BReC-CoV-2 which could diversify the immune response to mRNA which is normally dominated by B cells [60]. Deeper cellular analyses using these vaccines could help shed light on their unique mechanisms of protection. To further enhance the translational ability of this work, a model of waning immunity would also need to be established to mimic the changes in memory responses over time.

This VLP-RBD-BECC formulation shows promise as the COVID-19 pandemic progresses. Additional immunogenicity studies using Omicron RBD or other emerging strain RBDs will be necessary to improve the neutralizing antibody responses suggested by this challenge study. In these future immunogenicity studies, it will also be important to characterize the tissue-specific T cell responses to vaccination. Lung-resident T regulatory and T memory cells are vital to the lasting protection against SARS-CoV-2 [61–63]. Investigation of the neutralizing abilities of vaccine-induced IgA responses will also be interesting for describing not just the mechanistic ability of VLP-RBD-BECC to induce protection, but for also characterizing the potential of intranasal vaccines against other pathogens.

## Acknowledgments

We thank Drs. Laura Gibson and Clay Marsh of the West Virginia University Health Sciences Center for their support of COVID-19 preclinical research efforts. We would like to thank Drs. Matthew D.J. Dicks and Sumi Biswas (SpyBiotech) for the use of their VLP-RBD platform and Dr. Robert Ernst for the use of the BECC adjuvant. We thank Dr. Christopher Love (MIT) for providing the Spytagged RBD used in the VLP-RBD. We thank Dr. Ivan Martinez and Michael Winters for propagating the original challenge strain of the SARS-CoV-2 Delta variant, and Dr. Luis Martinez-Sobrido and Dr. Jordi Torrelles from the Texas Biomedical Research Institute for providing us with stocks of the Omicron variant for preclinical challenge studies. We would also like to acknowledge Mary Tomago-Chesney for her preparation of lung tissues for histology. We want to thank WVU’s Electron Microscopy, Histopathology and Tissue Bank Core for the images of lung tissues. The preparation of figures for this manuscript was supported by GraphPad Prism and BioRender.

## Author Contributions

These studies were designed by FHD, KSL, TYW and JRB. The BECC470 adjuvant was developed and provided by RKE. Viral strains were propagated and sequenced by MTW and IM. Mouse challenge studies as well as the subsequent necropsy and tissue processing were performed by KSL, OAM and MC. ELISAs were performed by NAR, OAM, and KSL. MC performed plaque assays, plaque reduction assays, and *in vitro* antibody neutralization assays. qRT-PCR of tissue viral RNA was performed by OAM. Luminex assays were performed by KSL. Histopathological analyses of lung tissues were carried out by MSA. The data in this manuscript was prepared by KSL and FHD. All authors contributed to the writing and revision of this manuscript.

